# Different behavioural profiles between invasive and native nudibranchs: means for invasion success?

**DOI:** 10.1101/2023.04.13.536773

**Authors:** Armando Macali, Sara Ferretti, Serena Scozzafava, Claudio Carere

## Abstract

Behaviour is predicted to be a primary determinant of the success of the invasion process during the early phases of colonization. Comparing sympatric invaders and native species may provide a good approach to unravel behavioural traits involved in an invasion process. In this study, we carried out an experimental simulation of the introduction and the establishment phase into a new environment and assessed the expression of activity, alertness and habituation in a non-indigenous Mediterranean population of the South African nudibranch *Godiva quadricolor* comparing its profiles with those of the sympatric native *Cratena peregrina* and *Caloria quatrefagesi*. Individuals of these three species were subjected to three behavioural tests: spontaneous activity, carried out in the introduction phase (immediately after sampling) and after a week of acclimatization; alert test, in which a potential threat was simulated by means of a tactile stimulus; habituation test, where the same alert test stimulus was repeated five times at thirty- minute intervals. Native nudibranch had repeatable traits, although with species differences perhaps related to their different ecological niches. The comparison with the invasive species highlighted its low repeatability in activity levels, suggesting higher plasticity, a strong tendency to locomotor activity, and a marked sensitization in the habituation test. Such traits could play an important and active role in the ongoing invasion process.

## Introduction

Biological invasions bloomed in the last century (Essl et al., 2011; Lewis et al., 2016) with enormous impacts and concerns, making crucial to understand the behavioural mechanisms enabling some species, while others not, to spread successfully over their natural geographic range. Behavioural and personality traits favouring the dispersal and establishment of new vital populations are predicted to be key factors in the selection process that affect the success or failure across the different stages of the introduction process and individual-level behavioural variation is particularly relevant in this respect (Chapple et al. 2012, Canestrelli et al. 2016). Whether an organism needs to adapt to changes into a completely novel landscape, the ability to trim adaptive traits quickly is essential (Tuomainen and Candolin, 2011). While selection may favor some behavioural phenotypes over others, the degree to which an organism can plasticly amend behaviour (behavioural plasticity) is also crucial in determining the outcome of an invasion (Chapple et al., 2012; Griffin et al., 2016, Cordeschi et al. 2022).

The Phenotypic Plasticity Hypothesis (Torchyk & Jeschke, 2018) suggests that alien species should display a higher degree of plasticity than natives (Richards et al., 2006), enabling the expression of more advantageous phenotypes, like the ability to access to novel resources in new environments (Grey & Jackson, 2012; Reisinger et al., 2017; Sol et al., 2002, 2005, 2011; Sol & Lefebvre, 2000), the expression of anti-predator behaviour (Hazlett et al., 2003; Reisinger et al., 2017) and flexible habitat choice (Grey & Jackson, 2012; Stroud et al., 2019). However, only recently research has started focusing on the role of behavioural traits and their plasticity in promoting invasion success at the individual and population levels (e.g., Damas-Moreira et al. 2019, Brand et al., 2021; Chapple et al., 2022; Cordeschi et al. 2022).

Animal personality, the consistent behavioural differences among individuals, is predicted to exert a pivotal contribution to the success (Sagata & Lester, 2009; Wolf & Weissing 2012; Chapple et al., 2012; Carere and Gherardi, 2012) and impact (Cote et al., 2010; Fogarty et al., 2011; Juette et al., 2014) of biological invasions. In general, exploration and boldness have been considered driving traits typical of alien species, especially in the early phases of the invasion process (Chapple et al., 2011, 2012; Juette et al., 2014; Griffin et al., 2016; Damas-Moreira et al., 2019, 2020). Novel invaders indeed represent a non-random subset of source population where behavioural polymorphisms and plasticity undergo founder events along the different colonization phases of the new environment (Shine et al., 2011; Canestrelli et al., 2016).

Individuals carrying correlated suites of traits (i.e., behavioural syndromes, Sih et al., 2012) may better succeed in the introduction into new environments, enhancing the chance to interact with transport vectors (I first invasion stage), establish a non-native population through the exploitation of local resources (II stage) and furtherly spread to new habitats expressing competitive advantages over native species (III stage) (Blackburn et al., 2011).

Comparative studies between invaders and sympatric, closely related, native species provide a good opportunity to detect functional traits and selective forces at play behind an invasion process (Holway and Suarez, 1999; Rehage & Sih, 2004; Rehage et al., 2005; Phillips and Suarez, 2012; Damas-Moreira et al., 2019; Parvulescu et al., 2021). The studies carried out so far suggest that invasive species do not behave consistently when compared to the natives (hymenoptera, Monceau et al., 2015; lizards, Damas-Moreira et al., 2019). Indeed, native species are likely to experience consistent selection pressure on behavioural traits, and repeatability across time is expected (Sih et al., 2012). Instead, a lack of clear syndromes and a low repeatability of behavioural traits, as found in some invasive species, may reflect higher levels of plasticity.

Since its emergence in the mid-20th century, invasion biology has matured into a productive research field addressing questions of fundamental and applied importance. Not only has the number of empirical studies increased through time, but also has the number of competing, overlapping and, in some cases, contradictory hypotheses about biological invasions (Enders et al., 2020; Richardson & Pysek, 2008). Despite marine invertebrates represent the most plentiful component of global biodiversity and the complexity of their nervous systems, as well as of their behaviours, are often as complex and variegated as those of vertebrates (Brembs 2013), so far behavioural studies on invaders mostly rely on terrestrial species. Of the 13,867 known established alien species worldwide, only 26% are associated with aquatic habitats (Cuthbert et al., 2021), with the Mediterranean hosting over 800 species, one of the most impacted marine environments worldwide (Zenetos & Galanidi, 2020). In this study, we carried out an experimental simulation of the introduction and establishment phases into a new environment, and repeatedly quantified spontaneous activity (including velocity and acceleration), alertness, and habituation to a tactile stimulus in the invasive *Godiva quadricolor*, a nudibranch species endemic to South Africa recently spread in the Mediterranean Sea, and in the native *Cratena peregrina* (Gmelin, 1791) and *Caloria quatrefagesi* (Vayssière, 1888).

## Material & Methods

### Study species

The South-African nudibranch *G. quadricolor* is an Indian Ocean native species which in the last century spread in different tropical and temperate areas (Cervera et al., 2010). In the Mediterranean Sea, *G. quadricolor* was likely unintentionally introduced as stowaways by commercial mollusk trade and ballast water and conspicuous populations have been reported from brackish waters and coastal lagoons used for aquacultures purpose (Willan, 2004; Cervera et al., 2010; Cattaneo-Vietti et al., 1990; Macali et al., 2013; Villani & Martinez, 1993; Furfaro et al., 2018; Kučić & Lanča, 2018).

We used Mediterranean native facelinid nudibranchs as ecologically relevant species for comparisons with the invasive nudibranch *G. quadricolor* (Barnard, 1927). *Cratena peregrina* (Gmelin, 1791) and *Caloria quatrefagesi* (Vayssière, 1888) are among the most common native nudibranchs in the upper littoral zone as well as in coastal lagoons and marinas, where the invasive populations have established (Cattaneo-Vietti e Chemello, 1991; Macali et al., 2013; Kučić & Lanča, 2018). Like the native species, *G. quadricolor* is diurnal, it conspicuously stands in the open, forming dense populations with high dispersal potential. It is likely that both the native and the invasive species have recent histories related to human activity; however, *C. peregrina* and *C. quatrefagesi* have not been successfully introduced outside of their native range, likewise the wider distribution of their typical prey (hydrozoan species of the genus *Eudendrium*).

### Sampling and housing

Individuals were sampled in Mediterranean coastal lagoons environments between April 2019 and July 2020. Considering the tendency to prey on other nudibranchs of *G. quadricolor*, to prevent any effect of this predatory pressure on the behavioural traits of the native species (Abrams, 2000), we selected Mediterranean species sampling sites in ecologically suitable areas where the presence of the non-native species was not ascertained. To do so, a total of 34 individuals of *C. quatrefagesi* and *C. peregrina* were collected from the Nassa Channel of Orbetello Lake (Italy, Tuscany; 42°25’59’’N-11°09’30’’E; 1m depth on rocky substrate) and 32 specimens of *G. quadricolor* from the Roman Channel of Sabaudia Lake (Italy, Latium; 41°14’50’’N-13°02’16’’E; 1–2 m depths on rocky-mud flats) (Fig.1). Given that both invasive and native species populations are established in homogenous artificial landscapes, it can be assumed that they have access to similar microhabitats.

**Figure 1.**
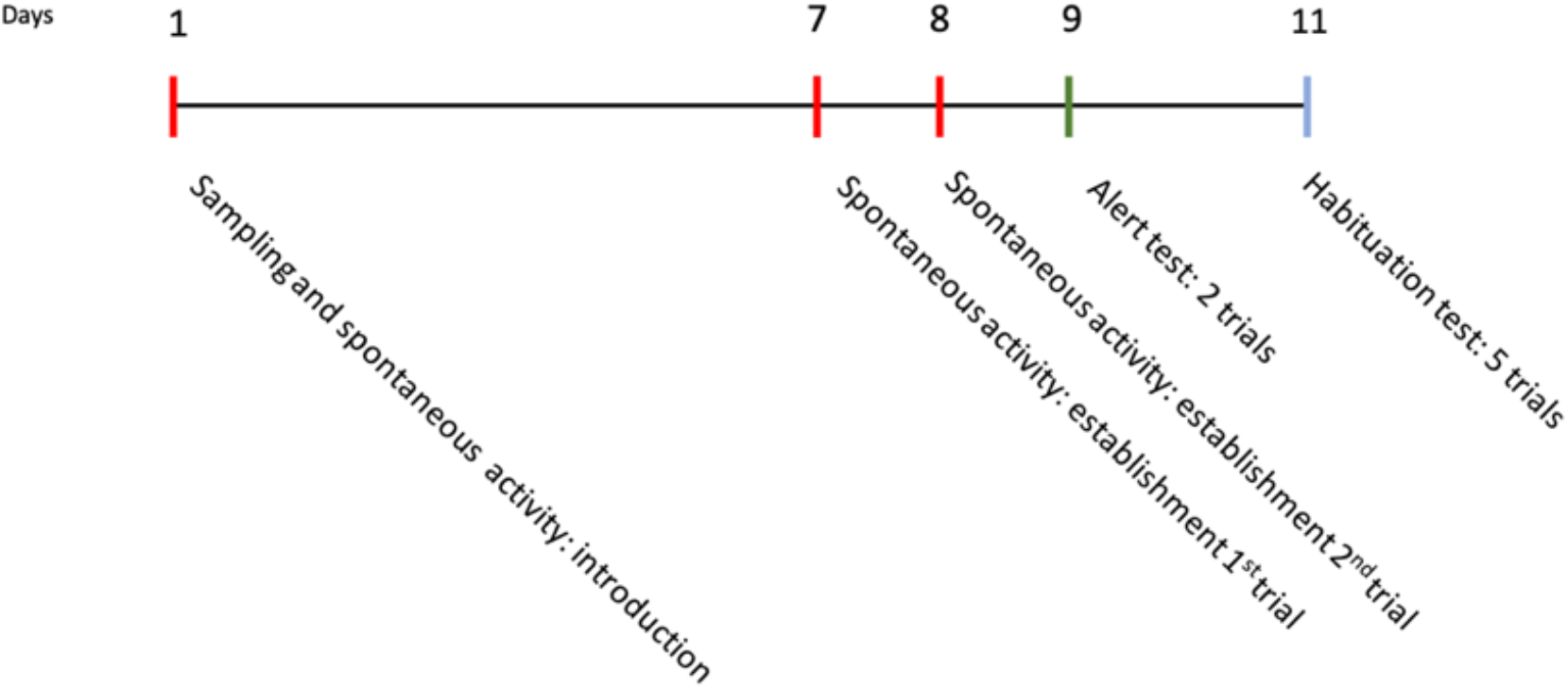
Schematic representation of the timeline of the behavioural tests to which the three nudibranch species were exposed. Red: activity test; green: alert test; light blue: habituation test.

All individuals were placed in separate plastic bottles filled with 0,5 l of marine water and transferred within two hours to the CISMAR (Marine Ichthyogenic Experimental Center, University of Tuscia) facilities. Each animal was photographed inside a Petri dish placed on graph paper and measured in length with the help of *ImageJ* software v 1.53. Animals were then randomly assigned to an experimental arena supplied with aerated running seawater at original environmental condition for the entire duration of the experiment (see Table 1 for details) and fed *at libitum* daily according to their specific diet, consisting of *Eudendrium sp*. for *C. quatrefagesi* and *C. peregrina*. As most of hunter opisthobranchs, *G. quadricolor* is considered a generalist species and euriphagy is well known also in other specialized carnivore nudibranchs (Megina et al., 2007); accordingly, animals were fed daily at libitum with *Artemia franciscana* and other nudibranch species.

**Table 1.** Behavioural scoring employed for each trial. Number (#) of individuals used for each trial: Cp: *Cratena peregrina*; Cq: *Caloria quatrefagesi*; Gq: *Godiva quadricolor*.

| Behaviour | Time | #Cp | #Cq | #Gq | Trait retrieved from video |
| --- | --- | --- | --- | --- | --- |
| **ACTIVITY**<br>*introduction (1 trial); establishment*<br>*(2 trials)* | 2 h | 30 | 20 | 21 | *Distance travelled (m)*<br>*Velocity (average, m/S)*<br>*Acceleration (average, M/S<sup>2</sup>)*<br>*Thigmotaxis: (% of path on external ring over the*<br>*whole path tracked)* |
| **CERATA EXTENSION**<br>**RESPONSE**<br>*Alert test* | 5 min | 29 | 15 | 13 | Time [s] of cerata extension after stimulus (2 trials,<br>12h interval) |
| **CERATA EXTENSION**<br>**RESPONSE**<br>*Habituation to alert test* | 5 min | 22 | 15 | 10 | *Time [s] of cerata extension after stimulus (5 trials,*<br>*30' intervals); occurrence of escape response.*<br>*Δt: difference in time (s) between the first and the*<br>*last trial in the habituation test.*<br>*Δe: difference in the frequency of escape behaviour*<br>*(e) between the first and the last trial in the*<br>*habituation test (e = escaping animals/N \*100)* |

### Behavioural assessment

The timeline of the behavioural tests is reported in Fig. 1

### Spontaneous activity

We measured spontaneous locomotory activity immediately after the disposal in the floating arenas, about three hours after the sampling, which simulates a sudden introduction into a new environment. The same measurements were taken one week later, which simulates an establishment. We considered two replicas of the test only for the establishment phase (7th and 8th day after the sampling) excluding the introduction trial to prevent any acclimatization biases in the test (Fig.1). Spontaneous activity was video recorded; given the differences in body length between species, we set up arenas proportionally to the average body size of the species, and grouped together, according to the different magnification settings required (n=16 for *G. quadricolor* and n=9 for *C. peregrina* and *C. quatrefagesi*). Videos were taken using a GoPro 5 set on time-lapse mode with a frame rate of 1f /5 sec for 120 minutes, with resolution of 1920×1080. The camera was mounted on a flexible “gooseneck”-style mounting arm for easy positioning in the center of the arenas block. Processing tracks to measure locomotory activity was performed with Tracker Video Analysis and Modeling Tools v5.1.5 (Open-Source Physics). Each time-lapse video was analyzed with a frame-by-frame method which refers to an interval of 5 second. Position of body control points was fixed on the upper part of the head due to its good visibility in each frame and replicability of selection. Before measurements, calibration was set according to the arena width. Motion analysis allowed determination of distance traveled, average velocity and average acceleration. We used the path track to quantify thigmotaxis, calculated as the percentage of path (as number of pixel) in the external ring (15 mm) of the arena over the whole path tracked.

### Cerata extension response: alertness and habituation test

Nudibranchs use cerata as a defensive tool against predators (Faulkner, 1992; Gavagnin *et al*., 1994; Aguado & Marin, 2007), displaying their active movement and extension in response to different stimuli under control of peripheral nerves and muscle contraction (Bickell-Page 1989). The individual cerata response to a tactile stimulus and the escape reaction (Video 1), was video recorded for 5 minutes right after the stimulus. The time elapsing between the extension of cerata upon the stimulus and their complete relaxation was measured as well as the occurrence of the escape response (Table 1).

Animals were individually transferred into smaller arenas, with 30 minutes of acclimatization before each test (see Tab. 1). We used a soft plastic string (10 × 3 × 0,1 mm) to gently apply the tactile stimulus in the median dorsal part of the animal, enough to induce cerata extension. We used this procedure in two different tests carried out on two consecutive days and differing in the time intervals between stimuli: in the *alert test*, two trials with an interval of 12h were conducted, whereas in the *habituation test*, five trials with an interval of 30 minutes between them were conducted (Fig.1).

### Statistical analysis

Statistical analyses were performed using SPSS, version 23.0 (SPSS Inc., Chicago, IL, USA) and RStudio, version 2022.07.1+554. We employed the RStudio ggplot2 package V.3.3.5 to graph spontaneous activity results.

We analysed the difference in size between the native and invasive species by means of a T-test for independent samples. In order to assess the contribution of size on the performance variables considered in the spontaneous activity, we used Pearson’s rank correlation test and simple linear regression between and within species.

We calculated individual-based repeatability across establishment trials in the animals’ response variables by intra-class correlation, calculated as the proportion of phenotypic variation that can be attributed to between-subject variation (Lessels and Boag, 1987). This was done by linear mixed-effects models (LMM) using individual identity as random factor for each species separately. The species differences in the response variables were tested by GLMMs. The correlations between behavioural traits for each species were further assessed using Pearson’s rank correlation tests.

## Results

### Repeatability

Repeatability of activity was higher in native species when compared to *G. quadricolor* (Fig. 2; SM Tab. 2). Among the native species, *C. quatrefagesi* showed lower repeatability, with speed and acceleration having p> 0.05, possibly due to the small sample size used in the study. (SM Tab. 2). Repeatability of alertness differed between native and invasive species: outputs from *C. peregrina* and *C. quatrefagesi* resulted significantly repeatable in the habituation test, whereas *G. quadricolor* showed significant coefficients only in the alertness test. Escape response was significantly repeatable for all species in both alert and habituation tests, except for *C. peregrina* in the alert test. Thigmotaxis was not significantly repeatable for any of the species (SM Tab. 2).

**Figure 2.**
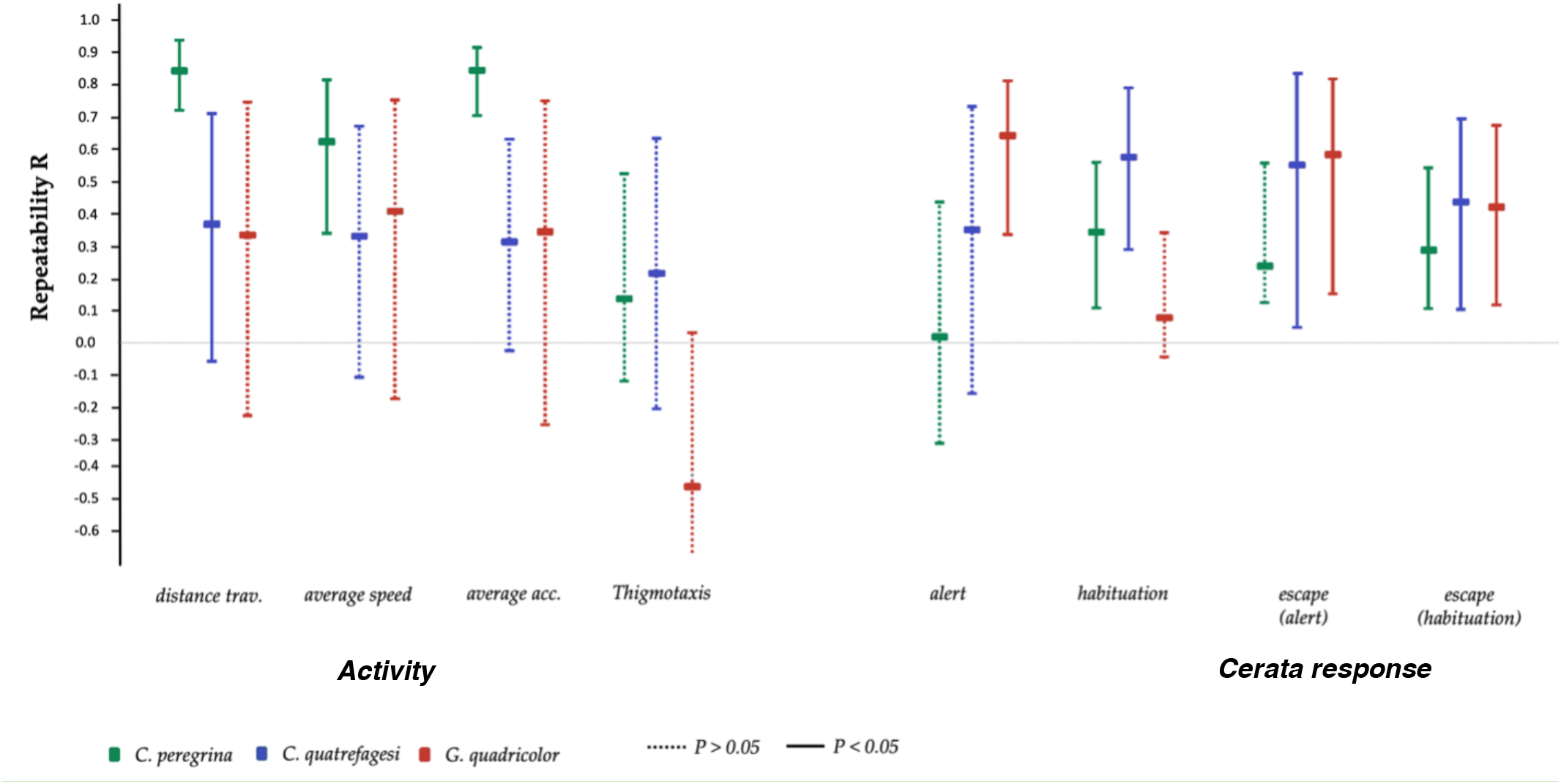
Repeatability (R) and 95% confidence intervals of behavioural traits (exploration and alertness) for the invasive *G. quadricolor* (red) and the natives *C. peregrina* (green) and *C. quatrefagesi* (blue).

**Table 2.**
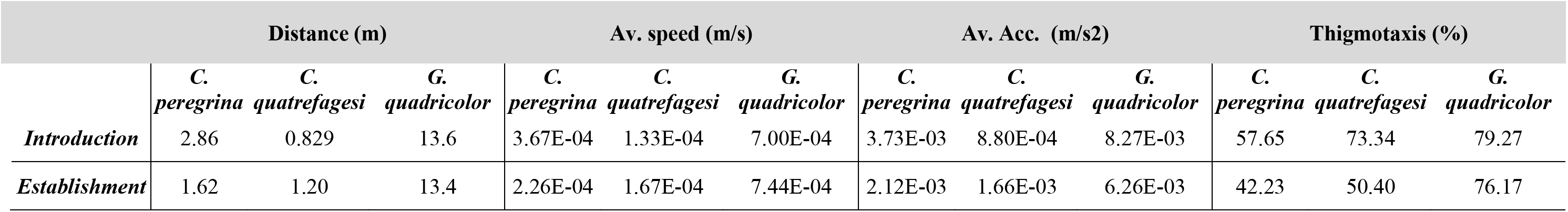
Mean value of activity trait retrieved in the native *C. peregrina, C. quatrefagesi* and the non-indigenous *G. quadricolor*

|  | Distance (m) |  |  | Av. speed (m/s) |  |  | Av. Acc. (m/s<sup>2</sup>) |  |  | Thigmotaxis (%) |  |  |
| --- | --- | --- | --- | --- | --- | --- | --- | --- | --- | --- | --- | --- |
|  | *C. peregrina* | *C. quatrefagesi* | *G. quadricolor* | *C. peregrina* | *C. quatrefagesi* | *G. quadricolor* | *C. peregrina* | *C. quatrefagesi* | *G. quadricolor* | *C. peregrina* | *C. quatrefagesi* | *G. quadricolor* |
| **Introduction** | 2.86 | 0.829 | 13.6 | 3.67E-04 | 1.33E-04 | 7.00E-04 | 3.73E-03 | 8.80E-04 | 8.27E-03 | 57.65 | 73.34 | 79.27 |
| **Establishment** | 1.62 | 1.20 | 13.4 | 2.26E-04 | 1.67E-04 | 7.44E-04 | 2.12E-03 | 1.66E-03 | 6.26E-03 | 42.23 | 50.40 | 76.17 |

### Correlations between traits

Overall, distance traveled and size were highly correlated with average speed, average acceleration and thigmotaxis, both in the introduction and the establishment phase (SM Tab. 3). Correlation between traits retrieved from the alertness and habituation test resulted significant only for habituation latency and Δt (the differences of latency of cerata extension between the first and the last trials), with differences between the introduction (R= 0.755; P<0.0001) and the establishment stage (R=-0.586; p<0.0001) (SM Tab. 3).

**Table 3.** Outcomes of the linear mixed-effects models for each exploratory behavioural trait. Significant results are in bold.

|  |  | Distance | Av. speed | Av. Acc. | Thigmotaxis |
| --- | --- | --- | --- | --- | --- |
| size | F | 7.447 | 6.584 | 8.948 | 1.08 |
|  | p | 0.007 | 0.011 | 0.003 | 0.3 |
| species | F | 35.909 | 10.250 | 5.146 | 4.474 |
|  | p | 0.001 | 0.001 | 0.007 | 0.013 |
| Phase | F | 1.842 | 0.876 | 4.268 | 0.251 |
|  | p | 0.177 | 0.351 | 0.041 | 0.617 |
| species*phase | F | 1.914 | 2.385 | 3.425 | 0.656 |
|  | p | 0.151 | 0.096 | 0.035 | 0.52 |

Correlations between traits within species differed significantly across experimental stages and species with only native species displaying a correlation between activity traits, both in the introduction and the establishment phase.

### Spontaneous activity

Overall, *G. quadricolor* was significantly more active than *C. peregrina* and *C. quatrefagesi* and the levels of activity increased as trials progressed; distance traveled, average velocity, acceleration and thigmotaxis were also higher when compared to native species but did not change as trials progressed (Tab. 2, Fig. 4).

The native species were significantly smaller in size (length) than *G. quadricolor* (t-test: *C. peregrina* and *G. quadricolor* F=11.423 p=0.001; *C. quatrefagesi* and *G. quadricolor* F=11.210 p=0.001).

Overall, distance travelled and size across species, were positively correlated in both the introduction and establishment phases (introduction R=0.769 p<0.0001; establishment R=0.865 p<0.0001). Within-species analysis (Fig. 3) revealed only a weak correlation in *C. peregrina* (R=0.357; p=0.038) in the introduction phase (*C. quatrefagesi* R=0.133; p=0.468; *G. quadricolor* R=0.280 p=0.312) and in *C. quatrefagesi* in the establishment phase (*C. quatrefagesi* R=0.473; p=0.026; *C. peregrina* R=0.056; p=0.763; *G*.*quadricolor* R=0.291 p=0.313).

**Fig. 3).**
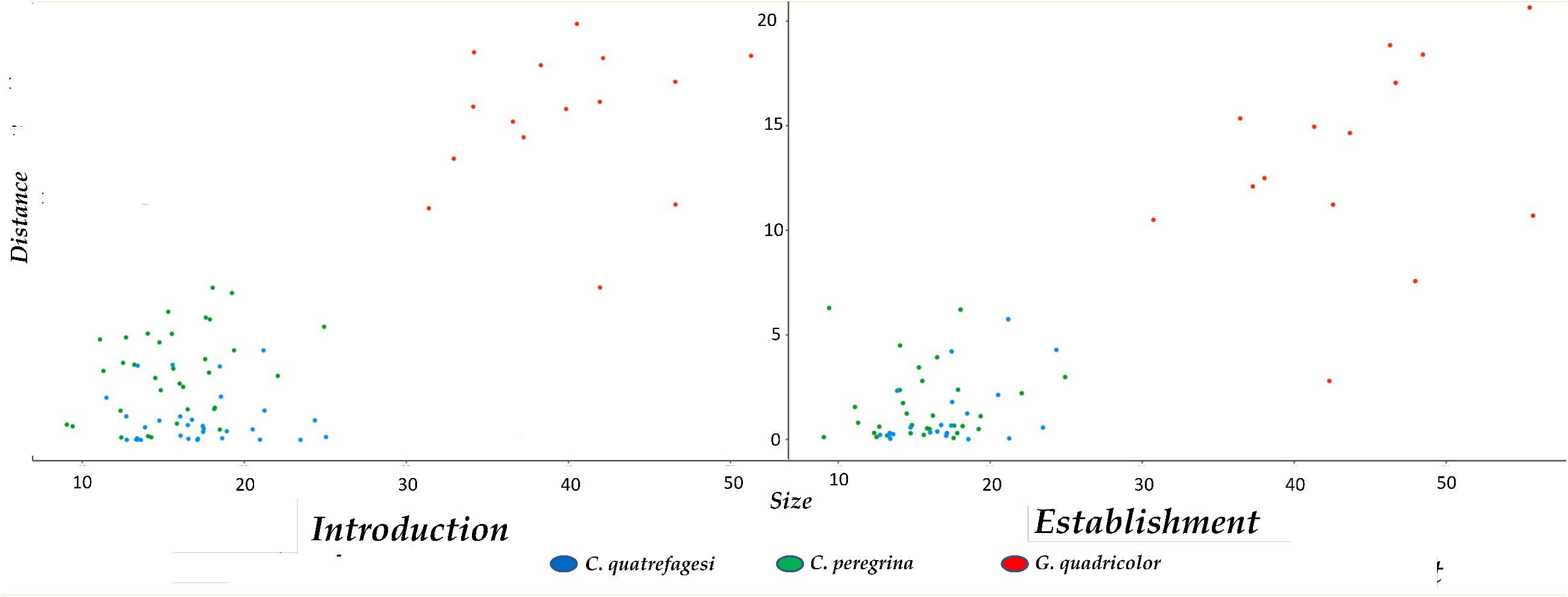
Simple linear regression graph between distance and size in the introduction and the establishment phase. In the introduction phase, 61.5 % of the distance variance is explained by size (Multiple R-squared: 0.6153) while in the establishment 74.7 % (Multiple R-squared: 0,7476).

**Figure 4.**
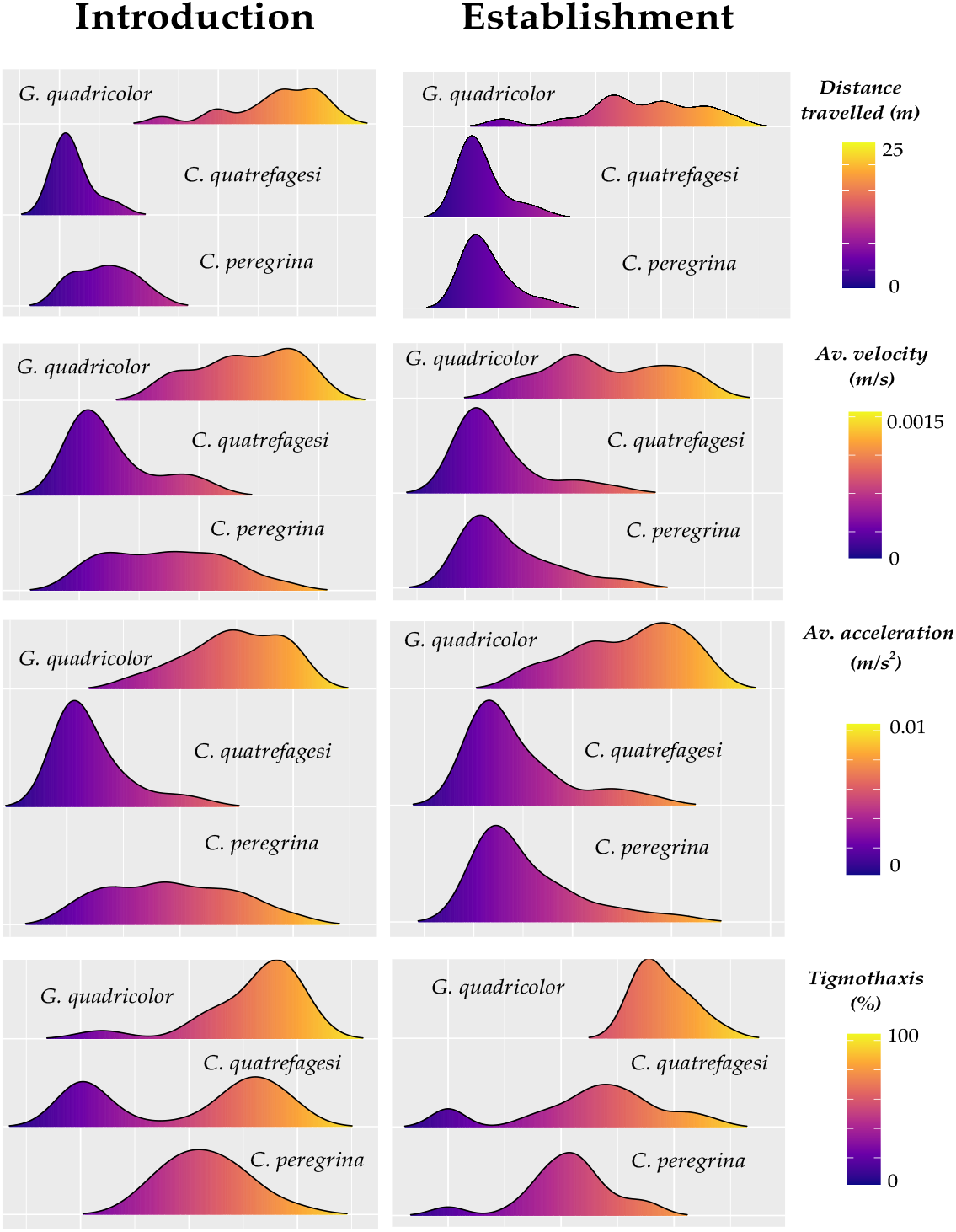
Kernel density plots shaping the distribution of the four behavioural traits, distance traveled, average velocity, average acceleration and thigmotaxis, measured in the three species during both the introduction and establishment tests.

### Cerata response: alertness and habituation

Differences in alertness were observed between native and alien species. *C. peregrina* and *C. quatrefagesi* both showed longer times of alertness as trials progressed whereas *G. quadricolor* quickly recovered from cerata extension in the second trial (Fig. 5). These results are supported by the significant interaction Trial*Species (Tab. 4).

**Figure 5.**
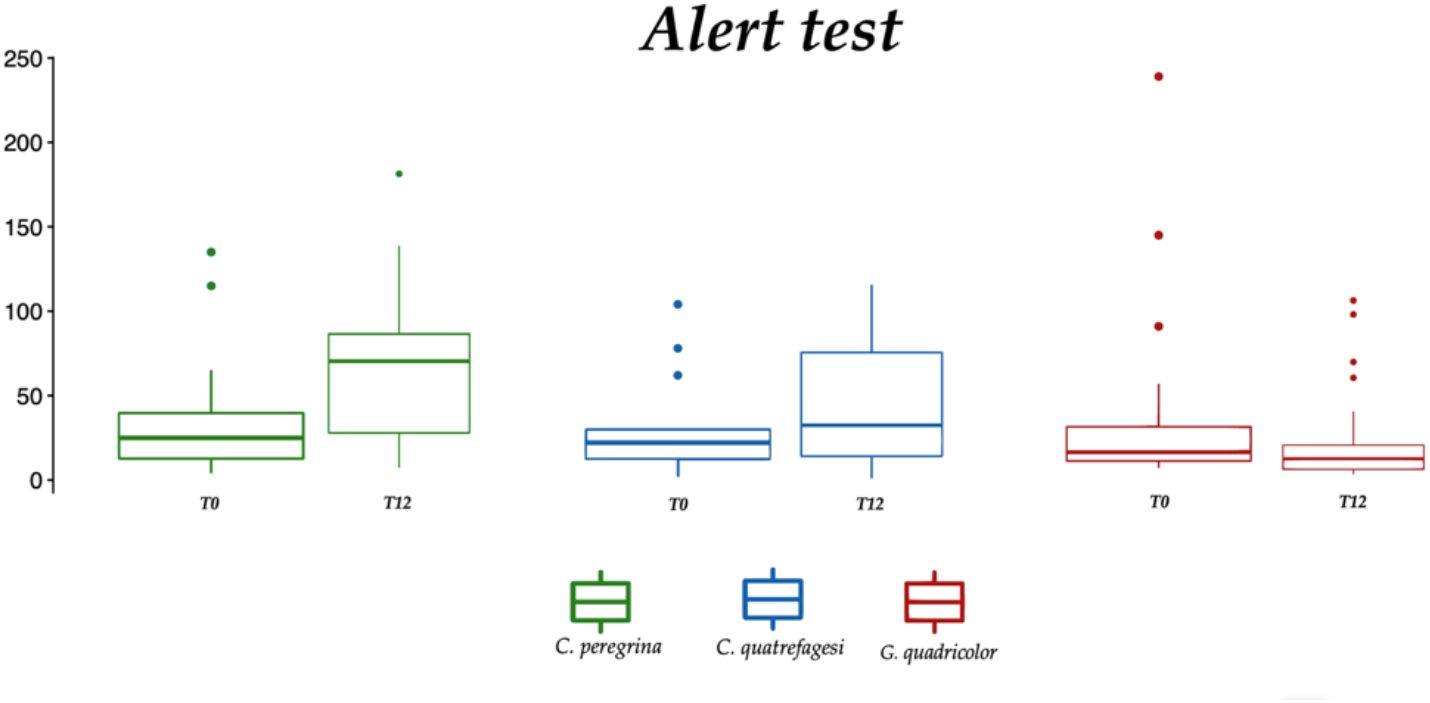
Boxplot of cerata extension time in alert test

**Table 4.**
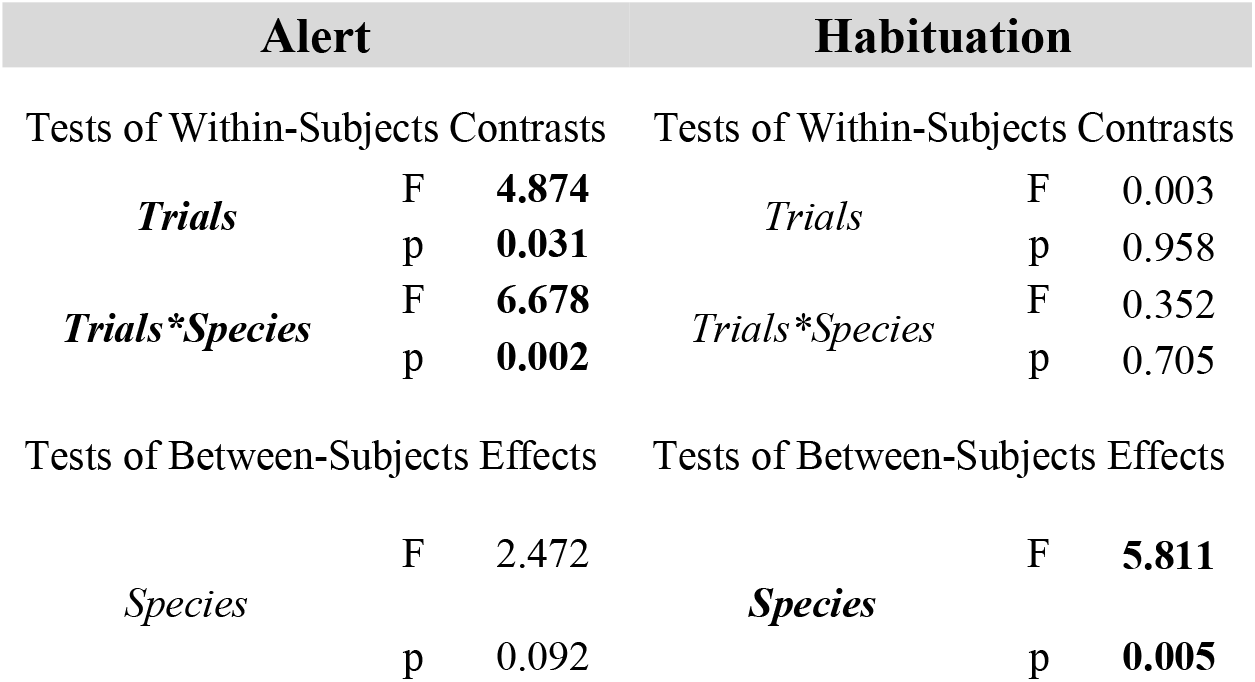
Outcome of the GLM for repeated measurements of cerata extension time after stimulus. Significant results are in bold.

In the habituation test, both native species showed a slight reduction in the time of cerata extension. On the contrary, *G. quadricolor* displayed an inverse profile, with a progressively increasing response, particularly marked between the third and fourth trial (Fig. 6a). Mean value of the ΔT, measured as the difference in cerata extension time between the first and the fifth trial. (Fig.6b) summarizes this pattern.

**Figure 6.**
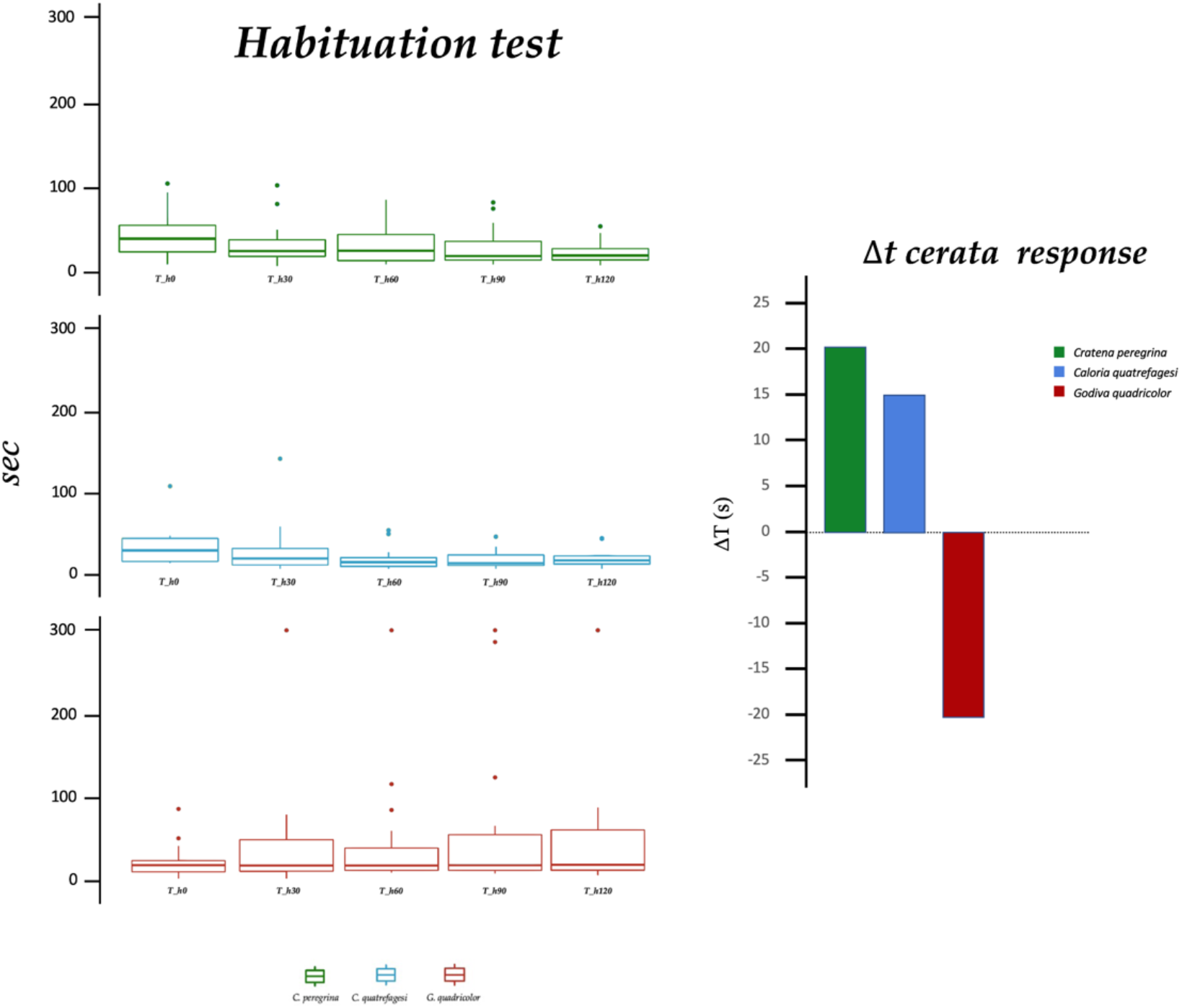
**a)** Boxplot of cerata expansion time in the habituation test; **b)**. ΔT is measured as the difference in cerata extension time between the first and the fifth trial.

Mean value of Δe, expressed as the difference in escape frequencies (escaping animals/N) between the first and last trial (Fig. 7), was inversely correlated with the frequency of stimulation in native species, suggesting reduced responsiveness to the stimulus over time. In contrast, *G. quadricolor* displayed a positive correlation between the frequency of stimulation and escape behaviour, with increased stimulation frequency leading to a stronger escape response.

**Figure 7.**
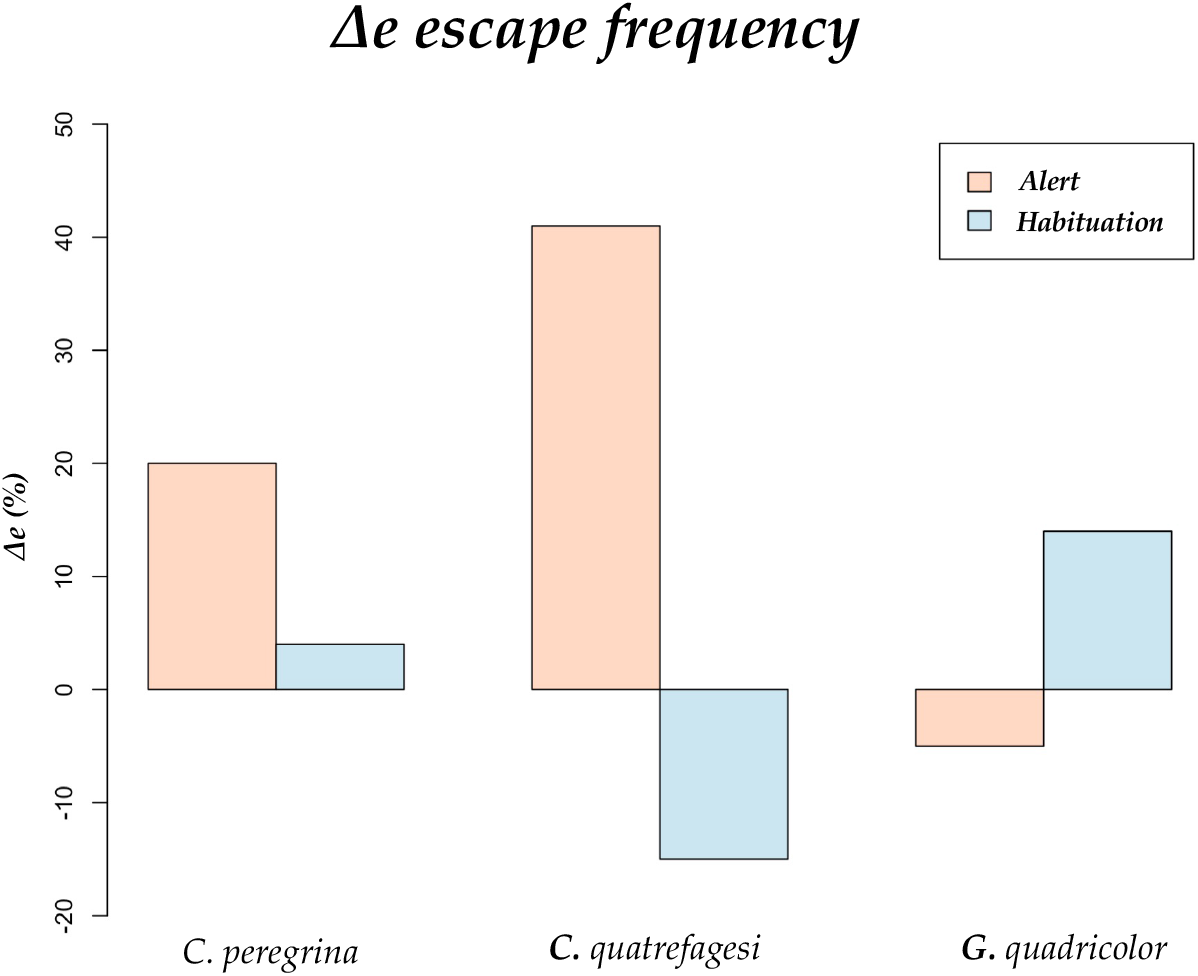
Mean value of the Δe, measured as the difference in escape frequencies (escaping animals/N) between the first and the fifth trial

## Discussion

Our comparative study illustrates clear differences in behavioural profiles and responsiveness between invasive and native nudibranch species. We report novel evidence in marine invertebrates suggesting a key role of behaviour in the spread of invasive species. Specifically, we found weak evidence for repeatability in activity profiles of the invasive species *G. quadricolor*, while the opposite emerged in the native *C. peregrina* and *C. quatrefagesi*. We argue that the reason for this outcome lies in the individual-level behavioural plasticity of the non-indigenous population, making behavioural choices in similar tests nonrepeatable. These findings fit with the higher behavioural plasticity reported in recent studies on non-indigenous species (Chapple et al., 2022; Damas-Moreira et al., 2019). Increased behavioural plasticity is thought to support organisms to cope with novel environmental, allowing them to rapidly adjust their behaviour to the new environmental conditions, particularly during the early stages of invasion, where invaders are characterized by small population sizes that are susceptible to environmental and demographic stochasticity (Chapple et al., 2022).

The repeatability of alertness in the nonindigenous species (Fig. 2) varied according to the frequency of the stimulation and the type of reaction, with outcomes symmetrically opposed to those retrieved from the natives for both experimental stages. The number of measurements is known to affect the overall repeatability, decreasing the measurement error associated with each observation, (Hoffmann 2000). Repeated measurement over a short timespan might also promote habituation to the stimuli, with individuals becoming less responsive or sensitized (Martin & Reale 2008). However, the lack of consistency of behaviours among individuals questions the possibility of detecting consistent behavioral differences in species having recently undergone a strong bottleneck or under new selective forces (Monceau et al., 2015).

*G. quadricolor* is a widespread invasive, frequently recorded in Mediterranean transitional waters and lagoons (Macali et al., 2013; Furfaro et al., 2018), with locally abundant population, up to 50 ind./100m 2 (Furfaro et al.,2018) and it is considered a generalist species. Conversely, *C. peregrina* and *C. quatrefagesi* occupy a more specialized niche, being selective predators on hydrozoan species. Despite the consistent size difference between natives and the invader, all three species display garish colors and a strong propensity to stand in the open. Colorful and vulnerable prey, such as most of the nudibranch species, are often protected against predation by using chemical defenses (Aguado & Marin, 2007) and, as for all aposematic species, advertised with conspicuous warning displays (Aguado & Marin, 2007; Ruxton et al., 2004). Although aposematism has been shown to reduce predation in several ways (Ruxton et al., 2004) aposematic prey do not always have lower predation than non-aposematic (Carroll & Sherratt, 2013; Seymoure et al., 2018; Yamazaki et al., 2020), resulting more subjected to repetitive and more cautious attacks if compared to its non-aposematic mutants (Sillén-Tullberg 1985; Yamazaki et al., 2020). Conspecific aggressiveness in hunter opisthobranchs is also known, with several species, including *G. quadricolor* but not *C. peregrina* and *C. quatrefagesi*, displaying cannibalistic behaviour (Macali et al., 2023). The repeatability of the alertness behaviour observed in *G. quadricolor* (Fig.2) may arise from new selective forces acting on the expression of antipredator strategies, especially in novel environments where they may be unfamiliar with local threats, at the same time, resulting exposed to high levels of cannibalism. Being more active, with greater dimension and different aposematic patterns, if compared to sympatric Mediterranean aeolids, *G. quadricolor* could have experienced increased predation pressure which may have promoted the sensitization of the cerata display (Fig. 6b) and a more stable expression of escape behaviours (Fig.2). The role of predator learning, indeed, is considered a driving force in the evolution of aposematic signaling (Barnet et al., 2007). In this framework, the enduring in the cerata extension along with the reduction of escaping behaviour as observed in the habituation test in *G. quadricolor* (Fig. 7) may thus be interpreted as a reinforcement of the aposematic signaling.

Consistent with the expected outlook, differences in activity levels between the invasive and native species were supported, with *G. quadricolor* traveling almost twice as far than natives (Fig. 4, Tab. 2). These results support previous studies showing that individuals on the expanding edge of the invasion front display enduring dispersal tendency and locomotor activity levels (Alford et al., 2009; Llewelyn et al., 2010).

The invasive *G. quadricolor*, displayed a strong propensity to thigmotactic behaviour during both the introduction and the establishment stage. Variation in the expression of thigmotaxis stems from different movement and perceptual strategies that influence the type and amount of information collected by animals (Doria et al., 2019). Thigmotaxis is commonly considered a measure of stress during captivity, hampering cognitive performance (Harris et al., 2009). Anxiety-related thigmotaxis usually decreases with increased familiarity with the experimental housing (Miller et al., 2018, Simon et al., 1994), as we observed in the native species. In contrast, we found weak differences in the proportion of time spent wall-following over the trials in *G. quadricolor*. While thigmotaxis has not been validated as a boldness assay, it is reasonable to assume a positive relationship between open space use and boldness (Carlson & Langkilde, 2013), with thigmotactic individuals reducing the risk of being detected by natural enemies (Harris et al., 2009). Correlations between exploration and other proactive behaviours have been underscored in different species (e.g., Evans et al., 2010; Scales et al., 2011; Canestrelli et al., 2016), drafting behavioural syndromes in which bolder individuals, or species, also tend to be more exploratory (Burstal et al., 2020; Chapple et al., 2011; Damas-Moreira et al., 2019; Monceau et al., 2015; Nordberg et al., 2021). In stable environments, proactive personality generally outcompetes reactive personality, this latter better adapted in changing environments (Koolhaas et al., 1999; Sih et al., 2004a, b). Under some circumstances, different trade-offs can be correlated to each behavioural type: proactive individuals, indeed, are frequently exposed to increased predation risk (Myers & Hyman, 2016; Nordberg et al., 2021), thus behavioural plasticity may be expected (Dingemanse, et al., 2010). The increased level of exploration activity along with a strong propensity to thigmotactic behaviour, as observed in *G. quadricolor*, may thus represent an effective tradeoff to reduce predation risk during the early phase of spread.

Behavioural traits of *C. peregrina* and *C. quatrefagesi* were similar, with exploratory individuals also being less sensitive to habituation in the alertness, suggesting a possible behavioural syndrome in this native species (Sih et al., 2004a, b). However, the same was not true for *G. quadricolor*, for which we did not find any relevant correlation between its behavioural traits. Nevertheless, the lack of correlation between traits, along with the overall inconsistency in *G. quadricolor* repeatability, may be advantageous during biological invasions. If variation in behavioural traits within a population increases the likelihood of the success of an invasion front (Dingemanse and Wolf, 2013, Sih et al., 2012), likewise, it constrains a population, since if selection acts on one trait, correlated behaviours are also likely to be affected (Sih et al., 2012).

In conclusion, our study sheds light on the role of behaviour in the ability to adapt to changing environmental conditions and contribute to their success in establishing new populations in an early stage of the invasion. By comparing the behavioural profiles of exploration and alertness in both native and invasive species, we were able to gain insights into the interplay between plasticity and personality and their impact on the success of the establishment of the South African nudibranch *G. quadricolor* in Mediterranean coastal environments. Whether these traits are the results of selective forces acting during the dispersal process or pre-existing plastic attributes of the species remains an open question. Additionally, we tested the use of anatomical structures, such as cerata, as a gauge for alertness in aposematic aeloids nudibranchs.

## Acknowledgments

The authors thank Giacomo Grignani and Alessandro Carlini for assistance during the experimental procedures. Bruno Bellisario gave statistical suggestions. The work was part of the Master theses in Ecology and Marine Biology of Serena Scozzafava and Sara Ferretti.

## Author Contributions

A.M. and C.C. designed research; A.M., S.F. and S.S performed research; A.M. and S.F. analyzed data with inputs from C.C.; A.M. wrote the paper with inputs from C.C. and S.F.

## References

Abrams PA, 2000. The evolution of predator-prey interactions: theory and evidence. Annu. Rev. Ecol. Evol. Syst. 31(1), 79–105.

Aguado F, Marin A, 2007. Warning coloration associated with nematocyst-based defences in aeolidiodean nudibranchs. J.Molluscan Stud 73(1): 23–28. https://doi.org/10.1093/mollus/eyl026

Alford RA, Brown GP, Schwarzkopf L, Phillips BL, Shine R, 2009. Comparisons through time and space suggest rapid evolution of dispersal behaviour in an invasive species. Wild Res 36(1):23–28.

Bickell-Page LR 1989. Autotomy of cerata by the nudibranch Melibe leonina (Mollusca): ultrastructure of the autotomy plane and neural correlate of the behaviour. Philos. Trans. R.Soc, 324(1222): 149–172. https://doi.org/10.1098/rstb.1989.0042

Blackburn TM, Pyšek P, Bacher S, Carlton JT, Duncan RP, Jarošík V, Richardson DM, 2011. A proposed unified framework for biological invasions. Trends Ecol. Evol 26(7): 333–339. https://doi.org/10.1016/j.tree.2011.03.023

Brand JA, Martin JM, Tan H, Mason RT, Orford JT, Hammer M P, Chapple DG, Wong BB, 2021. Rapid shifts in behavioural traits during a recent fish invasion. Behav. Ecol. Sociobiol., 75, 1–12.

Brembs B, 2013. Invertebrate behavior—actions or responses? Front. Neurosci 7:221. https://doi.org/10.3389/fnins.2013.00221

Burstal J, Clulow S, Colyvas K, Kark S, Griffin AS 2020. Radiotracking invasive spread: Are common mynas more active and exploratory on the invasion front? Biol. Invasions 22(8):2525–2543. https://doi.org/10.1007/s10530-020-02269-7

Canestrelli D, Bisconti R, Carere C 2016. Bolder takes all? The behavioral dimension of biogeography. Trends Ecol. Evol. 31(1): 35–43. https://doi.org/10.1016/j.tree.2015.11.004

Carere C, Gherardi F, 2012. Animal personalities matter for biological invasions. Trends Ecol. Evol, 28(1): 5–6. 10.1016/j.tree.2012.10.006

Carlson BE, Langkilde T, 2013. Personality traits are expressed in bullfrog tadpoles during open-field trials. J. Herpetol. 47(2):378–383.

Carroll J, Sherratt TN, 2013. A direct comparison of the effectiveness of two anti-predator strategies under field conditions. J. Zoo. 291(4):279–285.

Cattaneo-Vietti R, Chemello R, Giannuzzi-Savelli R, 1990. Atlas of Mediterranean nudibranchs. La Conchiglia., Rome, 264 pp.

Cattaneo-Vietti R, Chemello R, 1991. The opisthobranch fauna of a Mediterranean lagoon (Stagnone di Marsala, western Sicily). Malacologia 32: 291–299.

Cervera JL, Tamsouri N, Moukrim A, Villani G, 2010. New records of two alien opisthobranch molluscs from the north-eastern Atlantic: Polycera hedgpethi and Godiva quadricolor. Mar. Biodivers. Rec. 3(1). http://dx.doi.org/10.1017/S1755267210000102

Chapple DG, Simmonds SM, Wong BB, 2011. Know when to run, know when to hide: can behavioral differences explain the divergent invasion success of two sympatric lizards? Ecol. Evol 1 (3): 278–289. https://doi.org/10.1002/ece3.22

Chapple DG, Simmonds SM, Wong BB, 2012. Can behavioral and personality traits influence the success of unintentional species introductions? Trends Ecol. Evol. 27(1):57–64. https://doi.org/10.1016/j.tree.2011.09.010

Chapple DG, Naimo AC, Brand JA, Michelangeli M, Martin JM, Goulet CT, Wong B, 2022. Biological invasions as a selective filter driving behavioral divergence. Nat. Commun 13(1):1–11. https://doi.org/10.1038/s41467-022-33755-2.

Cuthbert RN, Pattison Z, Taylor NG, Verbrugge L, Diagne C, Ahmed DA, …, Courchamp F, 2021. Global economic costs of aquatic invasive alien species. Sci. Tot Environ 775, 145238.

Cordeschi G, Costantini D, Canestrelli D, 2022. Plastic Aliens: Developmental Plasticity and the Spread of Invasive Species. In Development Strategies and Biodiversity (pp. 267–282). Springer, Cham. https://doi.org/10.1007/978-3-030-90131-8_8

Cote J, Clobert J, Brodin T, Fogarty S, Sih A, 2010. Personality-dependent dispersal: characterization, ontogeny and consequences for spatially structured populations. Philos. Trans. R.Soc. Lond., B, Biol. Sci, 365(1560): 4065–4076. https://doi.org/10.1098/rstb.2010.0176

Damas-Moreira I, Riley JL, Carretero MA, Harris DJ, Whiting MJ, 2020. Getting ahead: exploitative competition by an invasive lizard. Behav. Ecol. and Sociobiol. 74(10): 1–12. https://doi.org/10.1007/s00265-020-02893-2

Damas-Moreira I, Riley JL, Harris DJ, Whiting MJ, 2019. Can behaviour explain invasion success? A comparison between sympatric invasive and native lizards. Anim. Behav. 151: 195–202. https://doi.org/10.1016/j.anbehav.2019.03.008

Dingemanse NJ, Kazem AJ, Réale D, Wright J, 2010. Behavioural reaction norms: animal personality meets individual plasticity. Trends Ecol. Evol. 25(2):81–89. https://doi.org/10.1016/j.tree.2009.07.013

Dingemanse N J, Wolf M, 2013. Between-individual differences in behavioural plasticity within populations: causes and consequences. Anim. Behav. 85(5), 1031–1039.

Doria MD, Morand-Ferron J, Bertram SM, 2019. Spatial cognitive performance is linked to thigmotaxis in field crickets. Anim. Behav. 150, 15–25.

Enders M, Havemann F, Ruland F, Bernard-Verdier M, Catford JA, Gomez-Aparicio L, Haider S, Heger T, Kueffer C, Kühn I, Meyerson LA, Musseau C, Novoa A, Ricciardi A, Sagouis A, Schittko C, Strayer DL,Vila M, Essl F, Jeschke JM, 2020. A conceptual map of invasion biology: Integrating hypotheses into a consensus network. Glob. Ecol. Biogeogr. 29(6):978e991. https://doi.org/10.1111/geb.13082

Essl F, Dullinger S, Rabitsch W, Hulme PE, Hülber K, Jarošik V, et al., 2011. Socioeconomic legacy yields an invasion debt. Proc. Nat. Acad. Sci. U.S.A. 108: 203–207. 10.1073/pnas.1011728108

Evans J, Boudreau K, Hyman J, 2010. Behavioural syndromes in urban and rural populations of song sparrows. Ethol. 116(7): 588–595. https://doi.org/10.1111/j.1439-0310.2010.01771.x

Faulkner DJ, Paul VJ, 1992. Chemical defences of marine molluscs., Ecological roles of marine natural products, Ithaca, New YorkComstock Publishing, pg. 119–163

Fisher RA, 1930. The genetical theory of natural selection. Oxford: Clarenden Press.

Fogarty S, Cote J, Sih A, 2011. Social personality polymorphism and the spread of invasive species: A model. Am. Nat. 177: 273–287. 10.1086/658174

Furfaro G, De Matteo S, Mariottini P, Giacobbe S, 2018. Ecological notes of the alien species Godiva quadricolor (Gastropoda: Nudibranchia) occurring in Faro Lake (Italy). J. Nat. Hist. 52(11-12): 645-657. http://dx.doi.org/10.1080/00222933.2018.1445788

Gavagnin M, Marin A, Castelluccio F, Villani G, Cimino G, 1994. Defensive relationships between Caulerpa prolifera and its shelled sacoglossan predators. J. Exp. Mar. Biol. Ecol. 175(2): 197–210. https://doi.org/10.1016/0022-0981(94)90026-4

Grey J, Jackson MC, 2012. ‘Leaves and eats shoots’: Direct terrestrial feeding can supplement invasive red swamp crayfish in times of need. PLoS ONE 7:e42575. https://doi.org/10.1371/journal.pone.0042575

Griffin AS, Guez D, Federspiel I, Diquelou M, Lermite F, 2016. Invading new environments: a mechanistic framework linking motor diversity and cognition to establishment success. Biol. Invasions Anim Behav. 26-46.

Harris AP, D’Eath RB, Healy SD, 2009. Environmental enrichment enhances spatial cognition in rats by reducing thigmotaxis (wall hugging) during testing. Anim. Behav. 77(6): 1459–1464.

Hazlett BA, Burba A, Gherardi F, Acquistapac P, 2003. Invasive species of crayfish use a broader range of predation-risk cues than native species. Biol. Invasions 5: 223–228. https://doi.org/10.1023/A:102611462361

Hoffmann A A, 2000. Laboratory and field heritabilities. Adaptive Genetic Variation in the Wild (TH Mousseau, B. Sinervo and JA Endler, eds.), 200–218

Holway DA, Suarez AV, 1999. Animal behavior: an essential component of invasion biology. Trends Ecol. Evol. 14(8): 328–330.

Juette T, Cucherousset J, Cote J, 2014. Animal personality and the ecological impacts of freshwater non-native species. Curr. Zool. 60(3):417–427. http://dx.doi.org/10.1093/czoolo/60.3.417

Koolhaas JM, Korte SM, De Boer SF, Van Der Vegt BJ, Van Reenen CG, Hopster H, Blokhuis HJ, 1999. Coping styles in animals: current status in behavior and stress-physiology. Neurosci Biobehav. Rev. 23(7):925–935. 10.1016/s0149-7634(99)00026-3.

Kučić M, Lanča L, 2018. First record of alien sea slug Godiva quadricolor (Barnard, 1927) (Gastropoda: Nudibranchia) in the eastern Adriatic Sea. Ann. Ser. Hist. Nat. 28:1-6. 10.19233/ASHN.2018.01

Lessels CM, Boag PT, 1987. Unrepeatable repeatabilities: a common mistake. Auk 104: 116–121. https://doi.org/10.2307/4087240

Lewis MA, Petrovskii SV, Potts JR, 2016. The Mathematics Behind Biological Invasions. (Switzerland: Springer).

Llewelyn J, Phillips BL, Alford RA, Schwarzkopf L, Shine R, 2010. Locomotor performance in an invasive species: cane toads from the invasion front have greater endurance, but not speed, compared to conspecifics from a long-colonised area. Oecologia 162(2): 343–348.

Macali A, Conde A, Smriglio C, Mariottini P, Crocetta F, 2013. The evolution of the molluscan biota of Sabaudia Lake: a matter of human history. Sci. Mar. 77(4):649–662. 10.3989/scimar.03858.05M

Macali A, Ferretti S, Scozzafava S, Carere C, 2023. Cannibalism, self-cannibalism and autotomy in the non-indigenous nudibranch Godiva quadricolor. Rend. Lincei Sci. Fis. Nat. (in press).

Martin JG, Réale D, 2008. Temperament, risk assessment and habituation to novelty in eastern chipmunks, Tamias striatus. Anim. Behav. 75(1), 309–318.

Megina C, Gosliner T, Cervera JL, 2007. The use of trophic resources by a generalist eolid nudibranch Hermissenda crassicornis (Mollusca: Gastropoda). Cah. Biol. Mar. 48(1), 1.

Miller AJ, Page RA, Bernal XE, 2018. Exploratory behavior of a native anuran species with high invasive potential. Anim. Cogn. 21(1): 55–65.

Monceau K, Moreau J, Poidatz J, Bonnard O, Thiéry D, 2015. Behavioral syndrome in a native and an invasive hymenoptera species. Insect Sci. 22(4):541–548.

Myers RE, Hyman J, 2016. Differences in measures of boldness even when underlying behavioral syndromes are present in two populations of the song sparrow (Melospiza melodia). J. Ethol. 34(3): 197–206. https://doi.org/10.1111/1744-7917.12140

Nordberg E, Denny R, Schwarzkopf L, 2021. Testing measures of boldness and exploratory activity in native versus invasive species: geckos as a model system. Anim. Behav. 177, 215–222.

Pârvulescu L, Stoia DI, Miok K, Ion MC, Puha AE, Sterie M, Aburel OM, 2021. Force and boldness: Cumulative assets of a successful crayfish invader. Front. Ecol. Evol. 9:49. https://doi.org/10.3389/fevo.2021.581247

Phillip BL, Suarez VS, 2012. The role of behavioral variation in the invasion of new areas. In: Candolin U, Wong BBM (eds) Behavioral responses to a changing world: mechanisms and consequences. Oxford University Press. pp. 190–200 DOI:10.1093/acprof:osobl/9780199602568.001.0001

Rehage JS, Sih A, 2004. Dispersal behavior, boldness, and the link to invasiveness: a comparison of four Gambusia species. Biol. Inv. 6(3):379–391. https://doi.org/10.1023/B:BINV.0000034618.93140.a5

Rehage JS, Barnett BK, Sih A, 2005. Foraging behaviour and invasiveness: do invasive Gambusia exhibit higher feeding rates and broader diets than their noninvasive relatives? Ecol. Freshw. Fish 14(4): 352–360. https://doi.org/10.1111/j.1600-0633.2005.00109.x

Reisinger LS, Elgin AK, Towle KM, Chan DJ, Lodge DM, 2017. The influence of evolution and plasticity on the behavior of an invasive crayfish. Biol. Inv. 19:815–830. https://doi.org/10.1007/s10530-016-1346-4

Richards CL, Bossdorf O, Muth NZ, Gurevitc J, Pigliucci M, 2006. Jack of all trades, master of some? On the role of phenotypic plasticity in plant invasions. Ecol. 9(8):981e993. https://doi.org/10.1111/j.1461-0248.2006.00950.x

Richardson DM, Pysek P, 2008. Fifty years of invasion ecology e the legacy of Charles Elton. Divers. Distrib. 14:161e168. https://doi.org/10.1111/j.1472-4642.2008.00464.x

Ruxton GD, Sherratt TN, Speed MP, 2004. Avoiding attack. The evolutionary ecology of crypsis, warning signals, and mimicry. Oxford University Press, oxford, 249 pp.

Sagata K and Lester PJ, 2009. Behavioral plasticity associated with propagule size, sources and the invasion success of the Argentine ant, Linepithema humile. J. Appl. Ecol. 46:19–27 https://doi.org/10.1111/j.1365-2664.2008.01523.x

Scales J, Hyman J, Hughes M, 2011. Behavioral syndromes break down in urban song sparrow populations. Ethol. 117(10):887–895. https://doi.org/10.1111/j.1439-0310.2011.01943.x

Seymoure BM, Raymundo A, McGraw KJ, Owen McMillan W, Rutowski RL, 2018. Environment-dependent attack rates of cryptic and aposematic butterflies. Curr. Zool. 64(5), 663–669.

Shine R, Brown GP, Phillips BL, 2011. An evolutionary process that assembles phenotypes through space rather than through time. PNAS 108(14):5708–5711. https://doi.org/10.1073/pnas.1018989108

Sih A, Bell A, Johnson JC, 2004a. Behavioral syndromes: an ecological and evolutionary overview. Trends. Ecol. Evol. 19(7):372–378. https://doi.org/10.1016/j.tree.2004.04.009

Sih A, Bell AM, Johnson JC, Ziemba RE, 2004b. Behavioral syndromes: an integrative overview. Q. Rev.Biol. 79(3):241–277. http://dx.doi.org/10.1086/422893

Sih A, Del Giudice M, 2012. Linking behavioural syndromes and cognition: a behavioural ecology perspective. Philos. Trans. R. Soc. 367:(1603), 2762–2772. https://doi.org/10.1098%2Frstb.2012.0216

Sillén-Tullberg B, 1985. Higher survival of an aposematic than of a cryptic form of a distasteful bug. Oecologia 67(3):411–415.

Simon P, Dupuis R, Costentin J, 1994. Thigmotaxis as an index of anxiety in mice. Influence of dopaminergic transmissions. Behav. Brain Res. 61(1):59–64.

Sol D, Duncan RP, Blackburn TM, Cassey P, Lefebvre L, 2005. Big brains, enhanced cognition, and response of birds to novel environments. PNAS 102:5460–5465. https://doi.org/10.1073/pnas.0408145102

Sol D, Lefebvre L, 2000. Behavioural flexibility predicts invasion success in birds introduced to New Zealand. Oikos 90:599–605. https://doi.org/10.1034/j.1600-0706.2000.900317.x

Sol D, Timmermans S, Lefebvre L, 2002. Behavioural flexibility and invasion success in birds. Anim. Behav. 63:495–502. https://doi.org/10.1006/anbe.2001.1953

Stroud JT, Colom M, Ferrer P, Palermo N, Vargas V, Cavallini M, Lopez J, Jones I, 2019. Behavioral shifts with urbanization may facilitate biological invasion of a widespread lizard. Urban Ecosyst. 22:425–434. https://doi.org/10.1007/s11252-019-0831-9

Torchyk O, Jeschke JM, 2018. Phenotypic plasticity hypothesis. In J. M. Jeschke, & T. Heger (Eds.), Invasion biology: Hypotheses and evidence (pp. 133e139). Wallingford, U.K: CABI. https://doi.org/10.1079/9781780647647.0000.

Tuomainen U, Candolin U, 2011. Behavioural responses to human-induced environmental change. Biol. Rev. 86:640–657. doi: 10.1111/j.1469-185X.2010.00164.x

Villani G, Martinez E, 1993. Some observations on the opisthobranch fauna from the Fusaro Lake, a brackish-water lagoon near Naples. Boll. Malacol. 29:201–209.

Yamazaki Y, Pagani-Núñez E, Sota T, Barnett CR, 2020. The truth is in the detail: predators attack aposematic prey with less aggression than other prey types. Biol. J. Linn. Soc.131(2):332–343

Willan RC, 2004. Godiva quadricolor (Barnard, 1927) (Nudibranchia: Facelinidae) spreads into southern Queensland. The Beagle: Records of the Museums and Art Galleries of the Northern Territory 20:31–36. https://doi.org/10.1371%2Fjournal.pone.0068900

Wolf M, Weissing FJ, 2012. Animal personalities: consequences for ecology and evolution. Trends Ecol. Evol. 27(8):452–461. http://dx.doi.org/10.1016/j.tree.2012.05.001

Zenetos A, Galanidi M, 2020. Mediterranean non-indigenous species at the start of the 2020s: recent changes. Mar. Biodiv. Rec. 13(1), 1–17.

