## Supplemental Tables 1-3 for "Different behavioural profiles between invasive and native nudibranchs: means for invasion success?"

| *Species* | *Size* | *SEM.* |
|---|---|---|
| *C. peregrina* | 15.694 (9.06-24.93) | 0.57 |
| *C. quatrefagesi* | 17.139 (11.49-25.04) | 0.58 |
| *G. quadricolor* | 41.763 (30.74-55.72) | 1.16 |

**Table 1** Average size (mm) of sampled *C. peregrina. F. quatrefagesi* and *G. quadricolor. SEM: Standard Error Mean*

| | *C. peregrina* | | | | *C. quatrefagesi* | | | | *G. quadricolor* | | | |
|---|---|---|---|---|---|---|---|---|---|---|---|---|
| **Trait** | **n** | **R** | **P** | **CI (95%)** | **n** | **R** | **P** | **CI (95%)** | **n** | **R** | **P** | **CI (95%)** |
| **Distance trav.** | 30 | **0.858** | **0.000** | **0.723-0.930** | 18 | **0.382** | **0.048** | **-0.075-0.706** | 13 | 0.337 | 0.119 | -0.238-0.737 |
| **Average speed** | 30 | **0.633** | **0.000** | **0.358-0.806** | 19 | 0.345 | 0.068 | -0.117-0.684 | 13 | 0.417 | 0.069 | -0.148-0.777 |
| **Average acc.** | 30 | **0.849** | **0.000** | **0.708-0.925** | 21 | 0.331 | 0.061 | -0.095;0.665 | 13 | 0.353 | 0.108 | -0.221-0.745 |
| Thigmotaxis | 31 | 0.187 | 0.153 | -0.174-0.504 | 20 | 0.244 | 0.143 | -0.211-0.612 | 13 | -0.492 | 0.963 | -0.812-0.055 |
| **Alert** | 26 | 0.071 | 0.363 | -0.319-0.440 | 15 | 0.374 | 0.077 | -0.152-0.735 | 26 | **0.642** | **0.001** | **0.345-0.822** |
| **Habituation** | 24 | **0.337** | **0.000** | **0.152-0.548** | 15 | **0.541** | **0.000** | **0.312-0.773** | 20 | 0.098 | 0.094 | -0.040-0.330 |

**Table 2.** Repeatability (R) of the behavioural traits measured across trials in the *establishment* phase in *C. peregrina, F. quatrefagesi* and *G. quadricolor*, assorted with their *P*-values (*P*) and confidence intervals CI (95%).

| **Overall traits correlation** | *Introduction* | | *Establishment* | |
|---|---|---|---|---|
| | **R** | **P** | **R** | **P** |
| **Dimension *vs* Distance** | 0.850 | **0.000*** | 0.865 | **0.000*** |
| **Dimension *vs* Av. Speed** | 0.606 | **0.000*** | 0.661 | **0.000*** |
| **Dimension *vs* Av. Acc.** | 0.522 | **0.000*** | 0.647 | **0.000*** |
| Dimension *vs* Thigmotaxis | 0.245 | 0.028 | 0.560 | **0.000*** |
| **Distance *vs* Av. Speed** | 0.811 | **0.000*** | 0.843 | **0.000*** |
| **Distance *vs* Average Acc.** | 0.534 | **0.000*** | 0.846 | **0.000*** |
| **Distance *vs* Thigmotaxis** | 0.355 | **0.001*** | 0.489 | **0.000*** |
| Distance *vs* Alert | -0.314 | 0.021 | - | - |
| Distance *vs* Habituation | - | - | 0.491 | **0.000*** |
| Distance *vs* Δt | - | - | 0.374 | 0.010 |
| **Av. speed *vs* Av. Acc.** | 0.669 | **0.000*** | 0.960 | **0.000*** |
| **Av. speed *vs* Thigmotaxis** | 0.414 | **0.000*** | 0.439 | **0.000*** |
| Av. Acc. *vs* Habituation | 0.701 | **0.000*** | 0.301 | 0.040 |
| Av. Acc. *vs* Thigmotaxis | 0.227 | 0.042 | 0.416 | **0.000*** |
| Av. Acc. *vs* Δt | 0.719 | **0.000*** | - | - |
| Alert *vs* Thigmotaxis | - | - | 0.447 | **0.002*** |
| **Habituation vs Δt** | 0.755 | **0.000*** | 0.586 | **0.000*** |

**Table 3.** Pearson rank order correlations (R) and confidence intervals (P) between behavioural traits for the overall dataset including the *introduction* and *establishment* phase simulation. Significant results are in bold.
